# Sleep Identification Enabled by Supervised Training Algorithms (SIESTA): An open-source platform for automatic sleep staging of rodent polysomnographic data

**DOI:** 10.1101/2020.07.06.186940

**Authors:** Carlos S. Caldart, Raymond E. A. Sanchez, Miriam Ben-Hamo, Asad I. Beck, Tenley A. Weil, Jazmine G. Perez, Franck Kalume, Bingni W. Brunton, Horacio O. de la Iglesia

**Affiliations:** Department of Biology, University of Washington, Seattle, WA; Graduate Program in Neuroscience, University of Washington, Seattle, WA; National Institute of Mental Health, Bethesda, MD; Center for Integrative Brain Research, Seattle Children’s Research Institute, Seattle, WA; Department of Neurological Surgery, University of Washington, Seattle WA; Department of Pharmacology, University of Washington, Seattle WA

## Abstract

The temporal distribution of sleep stages is critical for the study of sleep function, regulation, and disorders in higher vertebrates. This temporal distribution is typically determined polysomnographically. In laboratory rodents, scoring of electrocorticography (ECoG) and electromyography (EMG) recordings is usually performed manually, where 5-10 second epochs are categorized as one of three specific stages: wakefulness, rapid-eye-movement (REM) sleep and non-REM (NREM) sleep. This process is laborious, time-consuming, and particularly impractical for large experimental cohorts with recordings lasting longer than 24 hours.

To circumvent this problem, we developed an open-source Python toolkit, **Sleep Identification Enabled by Supervised Training Algorithms** (SIESTA), that automates the detection of these three main behavioral stages in mice. Our supervised machine learning algorithm extracts features from the ECoG and EMG signals, then automatically scores recordings with a hierarchical classifier based on Bagging Random Forest approaches. We evaluated this approach on data collected from wild-type mice housed under both normal and different lighting conditions, as well as from a mutant mouse line with abnormal sleep phenotypes. To validate its performance on test data, we compared SIESTA with manually scored data and obtained F_1_ scores of 0.92 for wakefulness, 0.81 for REM, and 0.93 for NREM.

SIESTA has a user-friendly interface that can be used without coding expertise. To our knowledge, this is the first time that such a strategy has been developed using all open-source and freely available resources, and our aim is that SIESTA becomes a useful tool that facilitates further research of sleep in rodent models.

## Introduction

Sleep is a highly adaptive process that plays an important role in numerous physiological functions. Sleep generally occurs in two stages: rapid eye movement (REM) and non-rapid eye movement (NREM). These stages are characterized by distinct electrophysiological signatures, and the temporal distribution of these stages during sleep defines a ‘sleep architecture’ that can have important consequences for critical physiological and behavioral processes including memory consolidation, mood and cognitive function.

The study of sleep architecture in the context of disease is an area of great interest. Sleep disorders are extremely common, impacting over 100 million individuals in the United States alone, and contribute to or are co-morbid with a wide range of neurological, metabolic and psychiatric conditions.^1^

Sleep disorders are usually diagnosed using polysomnography, which measures multiple physiological signals including electroencephalography (EEG), electrooculography (EOG), chin and leg electromyography (EMG), electrocardiography (ECG), breathing effort, oxygen saturation and airflow.^2^ However, most experts regard EEG and EMG as the essential signals for the classification of behavioral states as either wake, NREM or REM.^3^ The characteristics of each stage are currently standardized and defined by the American Academy of Sleep Medicine (AASM) Scoring Manual.^4^ In humans NREM can be further subdivided into four distinct stages (N1 to N3), but this distinction is lost or ignored in studies of rodent models, with very few exceptions.^5^

Despite this standardization, trained professionals exhibit an overall inter-scorer agreement of 82.6%, which decreases to around 70% for N1, N3^6^ and sleep spindles.^7^ Recent findings suggest that this variability can be largely attributed to epochs that could be legitimately be assigned to multiple sleep stages.^8^ Unlike human sleep data, there is currently no “gold standard” for the manual classification of rodent sleep stage^9^ and manual scoring of sleep recordings is often error-prone, time-consuming, and generally must be done offline.

Attempts to develop automated algorithms for sleep staging date back to 1969,^10^ and have incorporated a wide range of approaches with varying degrees of success.^11^ Previous protocols have been developed and validated using both human and rodent data,^12^ often with good results but low adoption by the field. One reason for this limited reach could be because many methods were developed and described assuming the user has expertise in mathematics and computer programming. While more user-friendly approaches have been developed,^13^ they are often validated only on data acquired from wild-type (WT) animals or under standard laboratory conditions, or require the purchase of a commercial license and give little to no information about the underlying algorithm or training data used to validate the approach.

Considering the state of the field, and specifically the value to the circadian biology community of long-term chronic polysomnographic recording in studying the role of the circadian clock in regulating sleep/wake cycles, we have developed Sleep Identification Enabled by Supervised Training Algorithms (SIESTA): an open-source, automated sleep stage classification software toolkit in Python.

We first generated a training dataset consisting of 20 days of ECoG and EMG recordings obtained from both WT and transgenic mice with disturbed sleep under multiple experimental lighting conditions. For every 10 seconds of recording, we extracted 54 total features from ECoG and EMG signals, and used these data to evaluate a battery of supervised machine learning algorithms. We selected the bagging classifier using random forest, which showed the highest performance as determined by the F_1_ score. We aimed to reach an accuracy of at least 83% across behavioral states, reflecting the average inter-scorer reliability reported by the AASM.^6^ When evaluated on test data withheld from the training set, this classifier performed well, achieving F_1_ scores of 0.92 for wakefulness, 0.81 for REM, and 0.93 for NREM.

SIESTA is an open-source platform developed in Python. Our user-friendly graphical interface provides an intuitive workflow from data selection to final scored output and includes an optional training module where users can both train SIESTA on their own data and contribute to our freely available database of training data.

## Material and methods

### Animals and Housing Conditions

All experiments with animals were performed in accordance with animal protocols approved by the Office of Animal Welfare at the University of Washington. Mice with a heterozygous deletion of the *Scn1a* gene (*Scn1a*^+/-^), which are a model for an intractable form of epilepsy called Dravet syndrome (DS) and are hereafter referred to as DS mice, were generated by targeted deletion of the last exon, encoding domain IV from the S3 to S6 segment and the entire C-terminal tail of Na_V_1.1 channel, as previously described.^14^ The mice used in this study were generated by crossing heterozygous mutant mice of C57BL/6 background with wild-type (WT) C57BL/6 mice (both males and females of each genotype), resulting in only WT or heterozygous *Scn1a* mutant offspring.15 The DS mouse has been previously described to have deficits in both the circadian and homeostatic regulation of sleep.^15–17^

### Electrocorticographic Recordings

Sleep was recorded as previously described^18^. Briefly, mice were anesthetized with isoflurane and placed into a stereotaxic device where isoflurane anesthesia continued throughout surgery. Each mouse was implanted with ECoG electrodes, consisting of dental screws (Pinnacle Technology, Lawrence, KS; No. 8209: 0.10-in.). A midline incision was made above the skull. Recording electrodes were screwed through cranial holes as follows: over the left frontal cortex (1.5 mm lateral and 2 mm anterior to bregma) and over the right parietal cortex (1.5 mm lateral and 2 mm posterior to bregma), a ground electrode was placed over the visual cortex (1.5 mm lateral and 4.0 mm posterior to bregma), and a reference electrode was placed over the cerebellum (1.5 mm lateral and 6.5 mm posterior to bregma). EMG signals were obtained by placing a pair of silver wires into the neck muscles. The screws were connected, through silver wires, to a common 6-pin connector compatible with the Pinnacle recording device. The screws and connector were fixed to the skull with dental cement. Mice were implanted at between 3 and 4 months of age to account for the long duration of circadian sleep experiments. After surgery, mice were housed in single recording cages under a 12:12 light-dark (LD) cycle. Mice had a recovery period of 1 week and were then fitted with a preamplifier and tether, and connected to the Pinnacle Technology recording system, where they were allowed 1 day to acclimate before recording started. The ECoG and EMG signals were sampled at 400 Hz with low-pass filters of 80 Hz and 100 Hz, respectively. All recordings are saved and exported for processing using the European Data Format (EDF), an open-source file format commonly used in polysomnographic recording.

### Behavioral Experiments and Environmental Conditions

Our recordings came from experiments maintained in a ventilated, light-tight room under either a 12:12 LD cycle with 200-lux intensity or DD. To determine the effect of abrupt LD phase shifts on the phase of the circadian rhythm of sleep, we used a delay and advance “jetlag” protocol. After mice were entrained to a 12:12 LD cycle, a delay jetlag was initiated by extending the light phase by 6 hours (i.e. delaying the time of lights off by 6 hours). The mice stayed under this new 12:12 LD cycle until they re-entrained to the new cycle. The advance jetlag was initiated by shortening the light phase by 6 hours (i.e. advancing the time of lights off by 6 hours). To assess the endogenous circadian period of sleep, we released mice into DD for at least 14 days after they had synchronized to a 12:12 LD cycle.

### Signal Processing and Feature Engineering

We used a total of 54 features extracted for each 10-second epoch, calculated from raw ECoG and EMG data. ECoG and EMG signals were prefiltered using a Hanning filter with a moving window of one minute.

From ECoG signals, we calculated power, energy and amplitude values for a series of spectral bands as described in previous studies of the ECoG and EMG signatures of sleep stages^19^. We defined the alpha band as 8-13 Hz, beta as 20-40 Hz, sigma as 11-15 Hz, gamma as 35-45 Hz,^20^ and a frequency band indicating the occurrence of sleep spindles as 12-14 Hz. Because there is little consensus on the definition of mouse theta in the literature, we incorporated several previously described definitions as features (4-12 Hz,^21^ 6-9 Hz,^22^ 5.5-8.5 Hz 19 7-10 Hz^21^). Next, we calculated three different power ratios between frequency bands. The first is the theta-delta ratio, which is commonly incorporated into heuristics used to discriminate sleep and wake states5. The other two have been used for sleep spindle detection^23^ and are defined as the ratio of power values of frequencies between 0.5-20 Hz to those between 0.5-50 Hz; and the ratio of those between 11-16 Hz to those between 0.5-40 Hz. Additional features included the 90% spectral edge, 50% spectral mean, mean and median amplitude, the root mean square, variance, skewness, and kurtosis of the signal. Finally, we incorporated the number of zero crossings, the peak-to-peak range, and the spectral entropy of the signal.

From the EMG signal, we calculated the amplitude, signal variance, skewness, kurtosis, root mean square amplitude and the spectral entropy. A list of all the features and their descriptions can be found in Supplementary Table 1.

### Software Development

All the codes were written and tested in Python 2.7 and 3.7.3, using Spyder 3.3.2 and Jupyter 5.7.2 on Windows, MacOS and Linux operating systems. The scikit.learn package version 0.20.1 was used for validating the machine learning algorithms and Kivy 1.10.1 for designing the graphical interface. We used Pandas 0.23.4, Pickle 3.0, Numpy 1.15.4 and Scipy 1.1.0 for data management and signal processing.

### Supervised Learning Algorithm Selection

Using our manually-scored sleep data, we tested the ability of several algorithms to identify each sleep stage. We used methods of one-step classification, wherein an epoch was classified as either wake, REM or NREM in a single step using one classifier; and two-level hierarchical classification, wherein one classifier was used to score an epoch as either wake or sleep, and a second classifier was used to further divide the bouts of sleep as either NREM or REM. The full list of classifiers tested is given in Table 1^24–36^.

**Table 1.** Inter- and intra-rater reliability metrics for manual scores used in training datasets

| Inter-Rater Reliability |  |  |  |  |
| --- | --- | --- | --- | --- |
|  | Total | Wake | NREM | REM |
| Fleiss Kappa [95% CI] | 0.75<br>[0.744-0.757] | 0.76<br>[0.758-0.770] | 0.73<br>[0.726-0.738] | 0.78<br>[0.773-0.786] |
| Agreement | 87.34% | 93.55% | 89.04% | 84.73% |
| # Epochs (% of Total) | 12608 | 5934 (47.07%) | 6047 (47.96%) | 627 (4.97%) |
| Intra-Rater Reliability |  |  |  |  |
| Rater | All | Wake | NREM | REM |
| A | 98.53% | 98.89% | 98.62% | 95.14% |
| B | 91.60% | 93.62% | 87.71% | 95.00% |
| C | 96.17% | 96.21% | 96.32% | 93.85% |
| D | 96.90% | 97.47% | 97.12% | 89.33% |
| Average | 95.80% | 96.55% | 94.94% | 93.33% |

*Note.* Cells under intra-rater reliability represent % agreement. NREM = non-rapid eye movement; REM = rapid eye movement.

Because the random forest algorithm is at the core of SIESTA, we provide a brief discussion of its implementation as follows. The random forest algorithm is an ensemble learning method that takes into account the classification results of multiple decision trees^27^. Although effective, decision trees often generalize poorly and underperform on classification problems with a large number of input variables.^31^ To compensate for this, random forest classifiers utilize a combination of decision trees such that each tree depends on a random, independently sampled vector of the dataset that has the same distribution for all decision trees. After several trees are created, the random forest algorithm “votes” on the most popular class. A margin function is used to determine by how much the average number of votes from decision trees for the right class exceeds the vote total for any of the other classes (e.g. wake, NREM or REM).^31^

SIESTA employs two other similarly structured ensemble learning methods called bagging and gradient boosting,^37^ using random forest as the base classifier for each. A bagging classifier^33^ is an ensemble meta-estimator method that creates random individual results by training each classifier on a random redistribution of the training dataset, and then aggregating their individual predictions (by a vote) to form a final prediction. This meta-estimator is typically used to reduce the variance of the core method (in this case Random Forest). Similarly, gradient boosting is an ensemble learning method that boosts the performance of “weak learners” such as decision trees using a loss function, which determines how well the model fits the training data.^33^

In the screening for all algorithms we performed 20-fold cross validation with shuffle and used F_1_ score to assess performance. The F_1_ score is the harmonic mean of the precision and recall of the test. Precision is given by the number of true positives (TP) divided by the sum of false positives and TP, and recall is calculated as the number of TPs divided by the sum of false negatives and TPs. Assessing test performance using F_1_ score compensates for the imbalance in the occurrences of wake, NREM and REM bouts, and as such is widely used in machine learning-based approaches to automatic sleep scoring.^11^ All results are shown as mean with standard deviation.

### Inter- and intra-scorer reliability

To calculate inter-scorer reliability, we randomly selected a 24-hour recording from our training dataset and compared the agreement between 4 experienced manual scorers using both percent agreement and Fleiss’ kappa.^38^ In accordance with previous reports^39^ of inter-scorer reliability for sleep scoring between multiple scorers, we set the following levels of agreement for evaluating ***κ***: 0-0.20 = slight agreement, 0.21-0.40 = fair agreement, 0.41-0.60 = moderate agreement, 0.61-0.80 = substantial agreement, 0.81-1.0 = near perfect agreement. ***κ*** values are reported as being within a 95% confidence interval (CI).

We then calculated intra-scorer reliability, or how consistently each manual scorer classified a single epoch as being the same state when presented with the epoch multiple times. We wrote a custom R script to randomly select 600 10-second epochs from the same 24-hour recording session, duplicate each of these epochs 5 times, randomly insert them into the original data file, and finally randomize the order of all epochs in the data file. This new file was then scored by 4 experienced manual scorers, and the score consistency between duplicate epochs was evaluated using percent agreement for each individual scorer.

### Training Dataset

The training dataset consisted of 20 total days of recording from 20 different mice, scored by 4 different experts blind to experimental conditions. Each recording was a day long (24 hours), under several different environmental lighting conditions including LD (12h light – 12h dark), Constant Darkness (DD), and in the days immediately following induced jet lag. These conditions covered the most common light cycle paradigms used in the study of behavioral circadian rhythms, with DD and jet lag particularly being known to induce changes in sleep timing and quantity.^40^ Data from these experiments were selected to make the dataset more generalizable to recordings obtained under diverse experimental conditions.

### Correlation Matrix and Dendrogram

We generated a correlation matrix to visualize the Pearson correlation coefficients calculated for pairs of features. We used single linkage clustering to generate the feature order.^41^ The dendrogram measures the pairwise distance between the features using a threshold and metric to divide the measurements called the cophenetic correlation coefficient. This allowed us to cluster the variables according to the average distance between each subset of merged features.

### Dimensionality Reduction using Sequential Feature Selection

Sequential Feature Selection belongs to a family of greedy search algorithms that are used to reduce an initial d-dimensional feature space (where d is the number of features extracted from the EDF file) to a k-dimensional subset (where k is the target number of features, and k < d). These methods select an initial size for the subset of features, and sequentially find the features^42^ that are most informative at this timestep. The process is repeated at the next timestep, choosing the next feature subset depending on the previously selected features and the metric of the classifier. In the forward sequential selector, a small feature subset is selected, and the algorithm adds a single feature at a time to the initial subset. This process is repeated until all features are added back in. For the backward sequential selector, the initial subset is of size d or near d, and the algorithm removes one feature at each timestep.

## Results

### Inter- and intra-scorer reliability

To evaluate the consistency of manual scoring and ensure the robustness of our training dataset, we first calculated measures for both inter-scorer and intra-scorer reliability. Four experienced manual scorers from our laboratory were asked to score a total of 12,608 10-second epochs as wake, NREM or REM, and inter-scorer reliability was determined using both Fleiss’ kappa and percent agreement (Table 1). Across all three stages, manual scorers displayed substantial Fleiss’ kappa agreement particularly in epochs scored as wake and REM, with NREM Fleiss’ kappa agreement being slightly lower (Table 2). Interestingly, wake and NREM showed the highest percent agreement, with the lowest for REM bouts (Table 1).

**Table 2.** Classification accuracy by algorithm and method.

| Algorithm | One-step | Hierarchical | Hierarchical + 2 hrs. |
| --- | --- | --- | --- |
| Logistic Regression | 0.78 (0.081) | 0.35 (0.074) | 0.86 (0.060) |
| Linear Discriminant Analysis | 0.77 (0.090) | 0.85 (0.077) | 0.85 (0.077) |
| K-nearest Neighbors Classifier | 0.76 (0.074) | 0.51 (0.009) | 0.84 (0.060) |
| Decision Tree Classifier | 0.72 (0.076) | 0.84 (0.043) | 0.84 (0.038) |
| Gaussian Naïve Bayes | 0.52 (0.18) | 0.35 (0.87) | 0.60 (0.149) |
| Passive Aggressive Classifier | 0.71 (0.079) | 0.48 (0.055) | 0.77 (0.0113) |
| Ridge Classifier | 0.70 (0.082) | 0.85 (0.077) | 0.85 (0.077) |
| Logistic Regression, cross validation | 0.79 (0.077) | 0.36 (0.086) | 0.86 (0.062) |
| Bernoulli Naïve Bayes | 0.60 (0.163) | 0.50 (0.032) | 0.75 (0.118) |
| Nearest Centroid | 0.54 (0.129) | 0.49 (0.013) | 0.67 (0.135) |
| Random Forest Classifier | 0.80 (0.061) | 0.89 (0.030) | 0.89 (0.028) |
| Ada Boost Classifier | 0.79 (0.066) | 0.88 (0.040) | 0.88 (0.040) |
| Bagging Classifier, Decision Tree | 0.79 (0.061) | 0.89 (0.030) | 0.88 (0.033) |
| Extra Trees Classifier | 0.80 (0.063) | 0.89 (0.031) | 0.89 (0.033) |
| Gradient Boosting Classifier | 0.83 (0.057) | 0.90 (0.035) | 0.90 (0.035) |
| Perceptron | 0.72 (0.112) | 0.49 (0.006) | 0.78 (0.077) |
| **Bagging Classifier, Random Forest** | **0.83 (0.060)** | **0.91 (0.032)** | **0.91 (0.032)** |
| Bagging Classifier, Extra Trees | 0.82 (0.062) | 0.90 (0.034) | 0.90 (0.032) |

*Note.* Values represent $F_1$ scores (with standard deviation in parentheses). Hierarchical + 2 hrs. methods refers to the method of adding two hours of manually scored data to the training set for the classifier. All data are unscaled.

Next, we evaluated intra-rater reliability using percent agreement. We found that on average our four manual scorers were highly consistent in their scoring across all three states (Table 2). This agreement was slightly higher for wake bouts, and slightly lower for NREM and REM bouts (Table 2).

### One-step classifier with complete dataset

Once we established consistency and high inter-rater reliability in our manual scoring, we examined which commonly used supervised learning algorithms performed with the highest accuracy in sleep stage classification when trained on our dataset (see Table 2 and Methods section for complete list). We tested each algorithm with the feature values both scaled and without scaling, as the performance of some algorithms changed depending on scaling; the results of both cases are displayed in Table 2 and Supplementary Table 2. Although several algorithms produced an F_1_ score that was better than chance using the one-step approach, GBC and BCRF performed the highest F_1_ score.

Because both GBC and BCRF do not require scaled data as a prerequisite to their use^31,35^ they did not show a change in performance whether the data were scaled or unscaled.

### Hierarchical classifier with complete dataset

With this approach, we observed improved detection of NREM and REM states using most of the algorithms we tested, with a F_1_ score of 0.907 ± 0.032 in the case of BCRF, an improvement from the F_1_ score of 0.827 ± 0.059 obtained using the one-step approach (Table 2). This improvement was not of equal magnitude in all three states scored. Table 2 and Supplemental Table 2 illustrate the performance of each classifier in first distinguishing sleep and wake, and second distinguishing between NREM and REM. The BCRF algorithm was the highest performing in terms of classification speed and accuracy (Sup. Table 3) and was subsequently selected as the classifier for SIESTA.

### Generalization of algorithm across genotypes and training datasets

We validated the performance of SIESTA using a test dataset excluded from the training dataset. This test data included WT and DS mice, manually scored by the same experts that labeled the original training dataset.

First, we tested SIESTA with only the WT data, training the algorithm with all but one WT mouse, using the scores from these new data as targets to validate the scoring method. This process was repeated 20 times, with both the one-step and hierarchical classifiers. Complete results of the scoring can be seen in Table 3. Finally, we added the first 2 hours of the manually scored data of the target mice to the dataset to reduce the interindividual variability of the individual mice in the training process,^48^ and then re-trained the BCRF in the hierarchical classifier with this new dataset. Using this approach, we increased classification performance for all three stages (Awake: 0.942 [±0.017] to 0.956 [±0.031]; NREM: 0.936 [±0.019] to 0.952 [±0.039]; REM: 0.811 [±0.036] to 0.844 [±0.054]).

**Table 3.** Sleep stage classification accuracy using subsets of the training data.

| Method | Wild Type Training/Wild Type Validation |  |  |  |
| --- | --- | --- | --- | --- |
|  | Wake | Sleep | NREM | REM |
| One-Step | 0.94 (0.015) |  | 0.90 (0.017) | 0.77 (0.038) |
| Hierarchical | 0.94 (0.017) | 0.92 (0.021) | 0.94 (0.019) | 0.81 (0.036) |
| Hierarchical + 2 hrs. | 0.96 (0.031) | 0.94 (0.038) | 0.95 (0.039) | 0.84 (0.054) |

| Dravet Syndrome Training/Dravet Syndrome Validation |  |  |  |  |
| --- | --- | --- | --- | --- |
| Method | Wake | Sleep | NREM | REM |
| One-Step | 0.85 (0.059) |  | 0.82 (0.094) | 0.60 (0.048) |
| Hierarchical | 0.85 (0.061) | 0.82 (0.115) | 0.91 (0.054) | 0.70 (0.084) |
| Hierarchical + 2 hrs. | 0.94 (0.013) | 0.93 (0.028) | 0.95 (0.016) | 0.81 (0.054) |

| Wild Type Training/Dravet Syndrome Validation |  |  |  |  |
| --- | --- | --- | --- | --- |
| Method | Wake | Sleep | NREM | REM |
| One-Step | 0.84 (0.091) |  | 0.86 (0.048) | 0.68 (0.081) |
| Hierarchical | 0.85 (0.097) | 0.88 (0.058) | 0.88 (0.072) | 0.75 (0.082) |
| Hierarchical + 2 hrs. | 0.88 (0.049) | 0.89 (0.030) | 0.89 (0.037) | 0.76 (0.061) |

*Note.* Cells contain F<sub>1</sub> accuracy scores (and standard deviation in parentheses). Hierarchical + 2 hrs. methods refers to the method of adding two hours of manually scored data to the algorithm. All data is unscaled. Wild type database n= 14, Dravet syndrome database n=6. NREM = non-rapid eye movement; REM = rapid eye movement.

These three different approaches (One-step classifier, hierarchical classifier and hierarchical classifier with the first 2 hours to the dataset) were repeated using a dataset consisting of only DS mice. Compared to WT data, these F_1_ scores were lower than for the WT database in the first two methods, but increased in the last method (Table 3). This discrepancy could be due to the known circadian rhythm and sleep disturbances present in DS mice.

Lastly, we attempted to use a training dataset consisting of only data from WT mice to classify data from DS mice. Once again, the best method was the hierarchical classifier using the first 2 hours of the target mice (Table 3). Based on these results, we concluded that the performance of the algorithm is best when the target data and the data in the training dataset are similar, and that adding scores for the first 2 hours of the target data increases the scores in all cases (Table 3).

Although scoring sleep in 10-second increments is common, this temporal resolution can result in the misclassification of boundary epochs, as sleep stage transitions can happen on rapid timescales. Therefore, we also tested the performance of the algorithm when tasked with scoring data in 5-second epochs. We normalized the database and validated the scores with a wild-type and a DS mouse. The F_1_ scores for these mice were similar to those obtained using 10-second epochs, with a score for REM of 0.75 in WT and 0.83 in DS using the One-step classifier; and 0.76 in WT and 0.85 in DS with the hierarchical classifier (Supplementary Table 5). These results validated the utility of this method even when changing the epoch size.

### Feature evaluation and reduction

Next, we decided to further characterize the features that we extracted from our raw data for training. Most features were empirically chosen based on features used to identify sleep stages according to the AASM, as well as previously developed methods for automatic sleep scoring in rodents. The aim of this feature characterization was to find a subset of the most important features for successful classification. Reducing the number of features used for scoring would allow for a faster and more flexible method that could work in a real-time scoring paradigm without significantly reducing classification accuracy.

We computed a correlation matrix to determine the extent of the similarity between features used in training, and found highly correlated features (r>0.9) that appeared to cluster primarily in two main groups: one with features describing the amplitude of the signal (in several frequency bands of both the ECoG signal and EMG signals), and a second cluster that included the relative power of all but the delta band of the ECoG signal, along with other features that describe the frequency domain of the signal (the spectral mean and spectral entropy of ECoG).

To further characterize the clustering and correlation of the features used for training, we constructed a dendrogram (Sup. Fig. 2). We found 3 clusters and 1 independent variable. The first cluster consisted primarily of variables related to the amplitude of the signals, similar to the correlation matrix. The second cluster contained a mix of the relative power and the energy of most frequency bands. The third cluster contained the fewest features and included the relative power and amplitude of the delta band and the spectral characteristics of the EMG. The only feature that did not belong to any cluster was the second ECoG index (‘ECOGrel2’, the ratio of the relative power of the 0.5-20 Hz band and the 0.5-50 Hz band), which has been previously identified as being useful in automated approaches to sleep scoring^49^.

After identifying subsets of features that were highly correlated, we next sought to find the optimal subset of features for classifying sleep stages. We used both Sequential Forward Selection (SFS) and Sequential Backward Selection (SBS) to identify the features that were the most critical for sleep stage identification. In both selection algorithms the inflection point (the number of features at which increasing the size of the feature subset does not significantly improve the performance of the classifier^50^) of the performance curve was around 6 features (Figure 2). Both algorithms were run with the one-step classifier using BCRF three times, each with cross-validation performed five times, to evaluate the robustness of the feature selection (Sup. Table 2). In most cases the same features were identified as being the most important for algorithm performance, and the inflection point of the performance curve always occurred at 6-8 features. The list of all features found by the sequential selection algorithm can be found in Sup. Fig. 1. The top 5 features consistently identified by both SFS and SBS as being most important were: ECoG Zero crossing, EMG Amplitude, ECoG Sleep Spindle Hanning window (listed as ‘ECoG Spindlehan’ in Fig. 1 and Sup. Table 1), ECoG Delta power and EMG Spectral Entropy. Most of the features identified have been already reported to be relevant for classifying sleep stages, either by being used to score sleep (ECoG Delta^51^) or being present mainly in certain stages (sleep spindles^52^).

**Figure 1.**
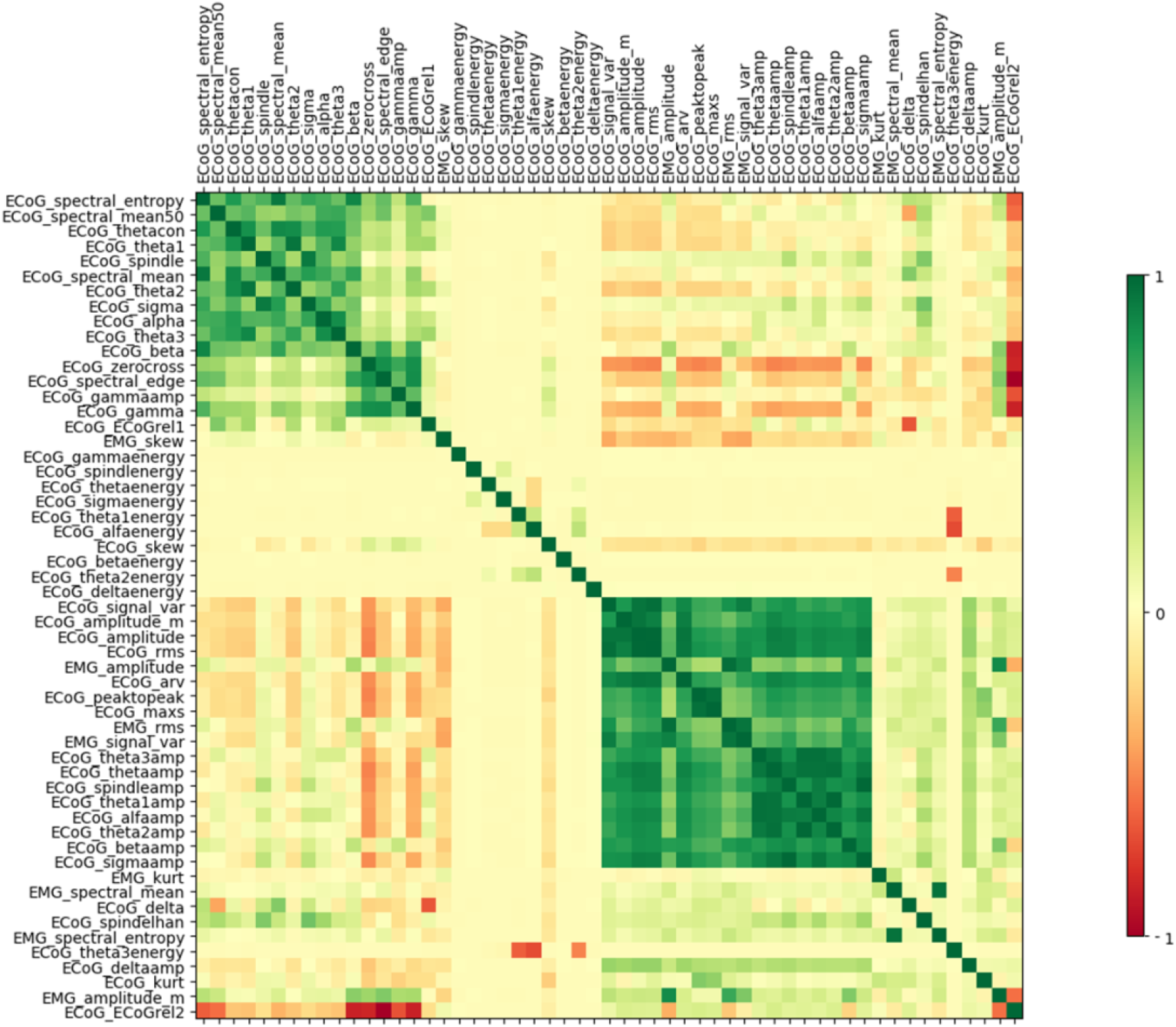
Correlation matrix of the features from the complete dataset. The data was reorganized via the clustering algorithm described in the Methods section to identify highly correlated subsets of features (in both positive – green - and negative – red - correlation). Each of the above features is described in detail in Supplementary Table 1.

**Figure 2.**
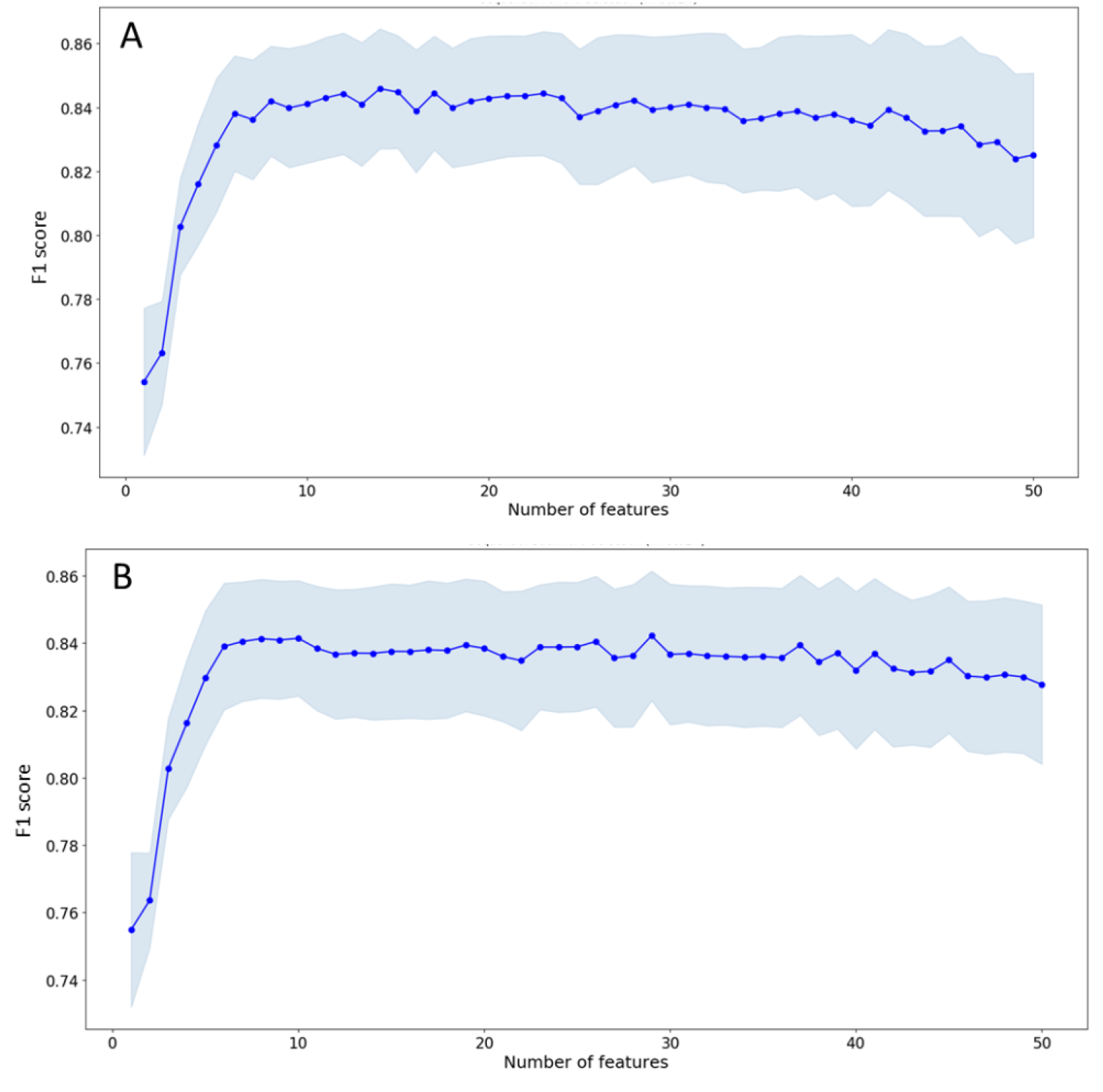
Performance under sequential feature selection of the BCRF algorithm trained on the complete training dataset using a One-Step approach to classification. The blue dots indicate the mean F_1_ values from 3 runs of the sequential feature selection method, and the light blue shaded area represents the standard deviation in each case. **A)** Sequential Forward Selection, **B)** Sequential Backward Selection.

As expected, several of the highest-rated features in all the iterations of the sequential search method were the same. We found 5 features that were present in all cases, and these features were spread across the three clusters identified in the dendrogram (Sup. Fig. 2) as well as the two most correlated clusters of the correlation matrix. Similarly, the correlation matrix revealed one feature identified by SBS and SFS in each cluster while the rest were not highly correlated with any other feature (Sup. Fig. 3).

### Graphical interface

Given the success of SIESTA in sleep stage classification, we sought to ease the access of our code to the broader sleep and circadian biology community. We developed a graphical interface that allows researchers to use our method, from pre-processing to scoring. SIESTA takes European Data Format (EDF) files as input, an open-source file format commonly used in both human and animal polysomnographic recordings. Each widget of the interface performs a different function, starting with either the feature extraction of new test data or training on a new dataset input by the user, to score new recordings. Users have the option to input parameters that are specific to their experimental needs, including the sampling frequency used in their recording and their desired epoch length. Both the features and final score files are output in .csv format. The stored database uses the Pickle library of Python to store the training results in a non-binary file so that users do not need to re-train the algorithm each time they want to score new data. A flow diagram of the complete training process and the actions available in the graphical interface is illustrated in Figure 3.

**Figure 3.**
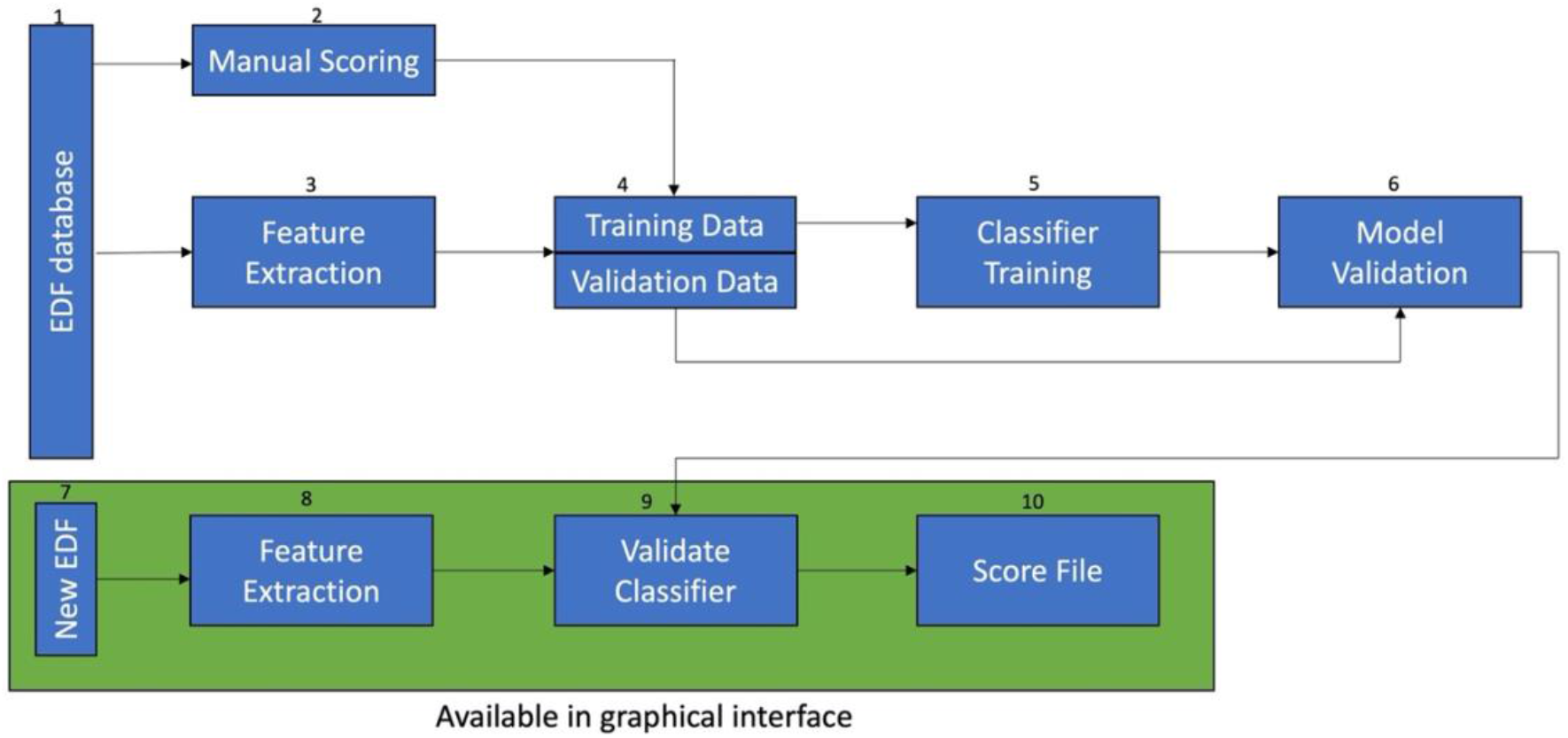
Overview of the SIESTA workflow. To train the model, we start with raw ECoG/EMG recordings in EDF file format **(1)**. Our dataset is manually scored by one of four human experts **(2)**. From data in the same EDF files, we calculated features used for scoring **(3).** Next, we split the data into training and validation sets **(4).** Using only the training set, we trained the BCRF classifier and obtain our model **(5).** We then evaluate model performance on the validation set **(6).** When a user wants to score data from their own experiment, they first upload their raw recording in EDF format **(7).** Feature extraction occurs as in Step 2, which generates a timestamped file containing feature values for every epoch in the recording. **(8).** Next, users can load the working model from our dataset. They will be informed of the validation results of the model as well as information about the underlying dataset (number and genotype of the animals included) **(9).** Users will also have the option to train a new classifier on their own manually scored dataset or combine their data with ours. Finally, users can upload the newly generated feature data to create a final output file containing timestamped scores and several metrics commonly used in sleep analysis, including ECoG delta power and theta power **(10).**

We freely provide the source code for the graphical interface, a user’s manual, and the database that we used to train our algorithm, as well as the base code that we used outside the graphical interface to validate SIESTA on GitHub^®^.

## Discussion

Using open-source coding and data management tools, we developed a novel automated sleep stage classification system for rodent polysomnographic data we call SIESTA. First, we compared the performance of a set of commonly used classifiers in identifying sleep stages from a training dataset containing data from both WT and transgenic mice with disturbed sleep. From this comparison, we identified the highest-performing classifier: the bagging classifier using random forest as the base classifier. We demonstrated that the accuracy of several classifiers was heavily influenced by the pre-processing of the signals, but the chosen method was not affected by this process. While other groups have reported using the random forest classifier to score sleep^43–45^ fewer have used the bagging^46^ or gradient boosting ensemble classification methods.^47^ To our knowledge, even fewer approaches have used the bagging classifier with random forest as the base classifier. We obtained high classification accuracy on par with AASM inter-rater reliability standards using a one-step classifier for all stages, and this score was further improved using a hierarchical classifier. We were able to improve classification accuracy even further when using a small amount (2 hours) of manually scored epochs from the data needing to be scored.

We have used this method to score data obtained in our laboratory, including continuous long-term polysomnographic recordings lasting over 3 weeks. Scoring this data manually is normally a tedious, error prone and time-consuming process. The run time for the feature extraction with the SIESTA code is less than a minute. In other words, we can score 24 hours of recording in under 1 minute if we use our complete training data set. If the algorithm is trained on new data, scoring 24 hours of recording takes approximately 5 minutes. Information on computational times associated with training and scoring with the different methods and subset of data can be found in Sup. Tables 3 and 4.

We welcome users to contribute both data and code to SIESTA to improve its performance and better suit their own needs. Accordingly, we provide the manually scored data we used to train the algorithm, as well as the source code of the classifier and feature extraction methods. We hope that as SIESTA is used in a wider range of experimental conditions and mouse lines, it will become more robust with a more comprehensive training dataset. A key feature of this code is that we do not use a time-dependent model, meaning the score of each epoch is independent from the previous epoch, so data from experimental animals with altered sleep architecture can be scored in an unbiased way. This characteristic allows for the training dataset to be replaced, modified or expanded with ease, allowing for more flexible scoring. Even though scoring can be improved when taking the previous epoch into account,^53^ this can lead to mistakes when the sleep architecture is atypical.

Scoring with SIESTA can be done without extensive coding knowledge, and our goal is that our simple interface and transparent analysis pipeline can make a novel contribution to the field of automatic sleep scoring and facilitate studies involving the chronic monitoring of sleep in the research community.^54^ Additionally, SIESTA dramatically reduces the time needed to obtain sleep scores from raw data compared to manual scoring. Our code is modular and separated into 3 main processes that comprise our analysis pipeline: feature extraction, training of the algorithm with the dataset and scoring of the experimental data file. All of the input and output files are in standardized open-source formats, making the manipulation of data using SIESTA easy. The motivation behind this open-source approach and the free release of SIESTA is to provide a trustworthy application that can be understood and validated by the community, encouraging any comment or modification that could improve the scoring process.

One of the main limitations of our approach is that we are not taking in account boundary epochs (epochs that include more than one state or a state change). These epochs can be difficult for both human and machine scorers to classify, and as such may decrease the efficiency of our algorithm. Additionally, neither our manual or automatic scorers have a criterion for the rejection of an epoch, so all data is stored in one of the predetermined labels (including epochs containing artifacts and transition epochs), which could also lower the performance of the algorithm. Even though we were aware of this limitation, we opted for this approach to simplify the use of the software and manual scoring of the recording. Even with this limitation, we obtain F_1_ scores of between 0.83 and 0.93 in all states.

Although we present only one pathological model in which we have validated our method, we are working on expanding the training dataset to include more data from additional mouse lines with altered sleep phenotypes. Additionally, we are looking to test the algorithm with data obtained from humans and other animal models, but there is recent work using similar approaches that lead us to believe our method can be effective in other models.^55,56^ Indeed, a recent study demonstrated that another supervised learning algorithm, deep convolutional neural networks, can be used to predict sleep stages from manually scored data in narcoleptic mice with a comparable degree of success as our own approach.^57^

An additional value of this work is the characterization of the features we use to train our model and score recordings. Through feature correlation and reduction, we specify a small subset of features needed to maintain high accuracy of SIESTA. Most of these features have been described in the literature to have a biological meaning (e.g. the relative power of the delta frequency band in the ECoG signal, EMG amplitude, and sleep spindles), which served to further validate our approach when they were found to be key in successful scoring. Other features are usually more prominently associated with other neural phenomena, as in the case of ECoG Zero-Crossing in the context of seizure and interictal spike detection.^58^ This might point to underlying biological processes that we have yet to fully characterize and are being identified by an unbiased machine learning algorithm. These supervised classifiers could also be finding success in scoring using the interaction between different features in the subset identified by SFS and SBS, as an emergent property of the complex ECoG and EMG features that by itself might not have a direct biological interpretation.^59,60^ Finally, the reduced subset of features we identified can be useful in the development of a real-time scoring paradigm, allowing more easily for long-term closed-loop manipulations of sleep stages in rodent models with both normal and pathological sleep, a potentially valuable tool for the sleep and circadian biology communities. One method that may increase availability of such real-time scoring is compressed sensing, which can reduce computational complexity and thereby computational time needed for sleep-scoring analyses^63^. This is effectively accomplished by only gathering data relevant to the task of interest, which tends to be a fraction of the overall data collected.

Although deep learning approaches have shown recent success in automated sleep scoring in both human patients^61^ and animal models^62^ (with large datasets and promising results), we opted for a shallow-learning approach without dimensionality reduction, allowing for greater interpretability of the results. The knowledge that the algorithm is using features with known biological relevance makes these results easier to approach and understand by the medical and research communities. Still, future studies should continue to directly compare the efficacy of supervised, unsupervised and deep learning approaches to scoring rodent polysomnographic data from diverse mouse lines and experimental conditions.

Finally, the study of the circadian regulation of sleep can produce long time series data that are often cumbersome to analyze. Our hope for SIESTA is not only to create an open-source community driven tool for sleep and circadian biologists, but also to encourage other researchers in the field not to shy away from performing experiments that would produce otherwise unwieldy datasets. These approaches are extremely valuable in furthering our understanding of the relationship between sleep, the circadian system and neurological and psychiatric conditions. Long-term monitoring of sleep in pre-clinical models of disease is an indispensable tool in understanding and leveraging these interactions to address the impact of sleep on health.

