## Supplementary Materials for "Sleep Identification Enabled by Supervised Training Algorithms (SIESTA): An open-source platform for automatic sleep staging of rodent polysomnographic data"

### This Document includes:

**Sup. Table 1 –** List of signal features used by SIESTA

**Sup. Table 2 –** Supervised learning algorithm validation using scaled training data

**Sup. Table 3 –** Run time for training each of the supervised learning algorithms validated on the complete training dataset

**Sup. Table 4 –** Run time for scoring with BCRF algorithm on different subsets of the data

**Sup. Table 5 –** SIESTA performance when scoring 5-second epochs

**Sup. Figure 1 –** Sequential feature selection performed using the one-step approach to classification

**Sup. Figure 2 –** Cluster dendrogram of the pair-wise distance of the features from the complete training dataset

**Sup. Figure 3 –** Correlation matrix with features identified by sequential feature selection highlighted

Supplementary Table 1 – List of Signal Features used by SIESTA

| | |
|---|---|
| **ECoG_delta** | Relative power of the 0.5-4 Hz frequency band |
| **ECoG_delta_energy** | Energy of the 0.5-4 Hz frequency band |
| **ECoG_delta_amp** | Mean amplitude of the signal in the 0.5-4 Hz frequency band |
| **ECoG_thetacon** | Relative power of the 4-12 Hz frequency band |
| **ECoG_thetaenergy** | Energy of the 4-12 Hz frequency band |
| **ECoG_thetaenergy** | Energy of the 4-12 Hz frequency band |
| **ECoG_thetamp** | Mean amplitude of the 4-12 Hz frequency band |
| **ECoG_theta1** | Relative power of the 6-9 Hz frequency band |
| **ECoG_theta1_energy** | Energy of the 6-9 Hz frequency band |
| **ECoG_theta1amp** | Mean amplitude of the 6-9 Hz frequency band |
| **ECoG_theta2** | Relative power of the 5.5-8.5 Hz frequency band |
| **ECoG_theta2energy** | Energy of the 5.5-8.5 Hz frequency band |
| **ECoG_theta2amp** | Mean amplitude of the 5.5-8.5 Hz frequency band |
| **ECoG_theta3** | Relative power of the 7-10 Hz frequency band |
| **ECoG_theta3energy** | Energy of the 7-10 Hz frequency band |
| **ECoG_theta3amp** | Mean amplitude of the 7-10 Hz frequency band |
| **ECoG_beta** | Relative power of the 20-40 Hz frequency band |
| **ECoG_betaenergy** | Energy of the 20-40 Hz frequency band |
| **ECoG_betamp** | Amplitude of the 20-40 Hz frequency band |
| **ECoG_alpha** | Relative power of the 8-13 Hz frequency band |
| **ECoG_alphaenergy** | Energy of the 8-13 Hz frequency band |
| **ECoG_alphaamp** | Mean amplitude of the 8-13 Hz frequency band |
| **ECoG_sigma** | Relative power of the 11-15 Hz frequency band |
| **ECoG_sigmaenergy** | Energy of the 11-15 Hz frequency band |

| | |
|---|---|
| **ECoG_sigmaamp** | Mean amplitude of the 11-15 Hz frequency band |
| **ECoG_spindle** | Relative power of the 12-14 Hz frequency band, the frequency at which NREM sleep spindles typically occur |
| **ECoG_spindlenergy** | Energy of the 12-14 Hz frequency band |
| **ECoG_spindleamp** | Mean amplitude of the 12-14 Hz frequency band |
| **ECoG_gamma** | Relative power of the 35-45 Hz frequency band |
| **ECoG_gammaenergy** | Energy of the 35-45 Hz frequency band |
| **ECoG_gammaamp** | Mean amplitude of the 35-45 Hz frequency band |
| **ECoG_ECoGrel1** | Ratio of the relative power value of the thetacon band (4-12 Hz) to that of the delta band (0.5-4 Hz) |
| **ECoG_ECoGrel2** | Ratio of the relative power value of the 0.5-20 Hz band to that of the 0.5-50 Hz band |
| **ECoG_Spindlehan** | Ratio of 11-16 Hz power to 0.5-40 Hz power smoothed with a 12-point Hanning filter |
| **ECoG_spectral_edge** | 90% spectral edge of the ECoG signal |
| **ECoG_spectral_mean50** | 50% spectral mean of the ECoG signal |
| **ECoG_zerocross** | Counts of the number of times the amplitude of the ECoG signal falls above or below the mean of the ECoG signal for any given epoch |
| **ECoG_maxs** | Maximum amplitude of the raw ECoG signal |
| **ECoG_peaktopeak** | Peak-to-peak amplitude of the ECoG signal |
| **ECoG_arv** | Arithmetic mean of the absolute values of the ECoG signal in a given epoch |
| **ECoG_rms** | Root mean square value of the ECoG signal |
| **ECoG_amplitude** | Mean amplitude of the ECoG signal in a given epoch |
| **ECoG_amplitude_m** | Median amplitude of the ECoG signal in a given epoch |
| **ECoG_signal_var** | Spectral variance of the ECoG signal |

| | |
|---|---|
| **ECoG_skew** | Skewness of the ECoG signal |
| **ECoG_kurt** | Kurtosis of the ECoG signal |
| **ECoG_spectral_mean** | Mean of the spectral power distribution of the ECoG signal for a given epoch |
| **ECoG_spectral_entropy** | Entropy of the spectral power distribution of the ECoG signal for a given epoch |
| **EMG_amplitude** | Mean amplitude of the EMG signal |
| **EMG_signal_var** | Spectral variance of the EMG signal |
| **EMG_skew** | Skewness of the EMG signal |
| **EMG_kurt** | Kurtosis of the EMG signal |
| **EMG_spectral_mean** | Mean of the spectral power distribution of the EMG signal for a given epoch |
| **EMG_spectral_entropy** | Entropy of the spectral power distribution of the EMG signal for a given epoch |
| **EMG_amplitude_m** | Median amplitude of the EMG signal in a given epoch |

Supplementary Table 2. Classification accuracy by algorithm and method when data are scaled.

| Algorithm | One-step | Hierarchical | Hierarchical + 2 hrs. |
|---|---|---|---|
| Logistic Regression | 0.80 (0.067) | 0.47 (0.005) | 0.87 (0.054) |
| Linear Discriminant Analysis | 0.77 (0.090) | 0.86 (0.056) | 0.86 (0.056) |
| K-nearest Neighbors Classifier | 0.76 (0.075) | 0.48 (0.003) | 0.85 (0.050) |
| Decision Tree Classifier | 0.72 (0.078) | 0.80 (0.061) | 0.80 (0.063) |
| Gaussian Naïve Bayes | 0.52 (0.18) | 0.48 (0.005) | 0.79 (0.136) |
| Passive Aggressive Classifier | 0.68 (0.11) | 0.46 (0.033) | 0.84 (0.052) |
| Ridge Classifier | 0.70 (0.083) | 0.78 (0.072) | 0.78 (0.072) |
| Logistic Regression, cross validation | 0.79 (0.077) | 0.48 (0.005) | 0.87 (0.062) |
| Bernoulli Naïve Bayes | 0.60 (0.163) | 0.48 (0.009) | 0.69 (0.123) |
| Nearest Centroid | 0.54 (0.129) | 0.39 (0.012) | 0.79 (0.125) |
| Random Forest Classifier | 0.80 (0.063) | 0.86 (0.051) | 0.86 (0.051) |
| Ada Boost Classifier | 0.79 (0.066) | 0.86 (0.05) | 0.86 (0.050) |
| Bagging Classifier, Decision Tree | 0.79 (0.06) | 0.85 (0.054) | 0.85 (0.054) |
| Extra Trees Classifier | 0.80 (0.062) | 0.86 (0.053) | 0.86 (0.053) |
| Gradient Boosting Classifier | 0.83 (0.057) | 0.86 (0.051) | 0.86 (0.051) |
| Perceptron | 0.72 (0.11) | 0.40 (0.01) | 0.84 (0.057) |
| **Bagging Classifier, Random Forest** | **0.83 (0.060)** | **0.87 (0.055)** | **0.87 (0.055)** |
| Bagging Classifier, Extra Trees | 0.82 (0.061) | 0.87 (0.052) | 0.87 (0.054) |

*Note*. Values represent $F_1$ scores (with standard deviation in parentheses). Hierarchical + 2 hrs. methods refers to the method of adding two hours of manually scored data to the training set for the classifier. All data are scaled.

Supplementary Table 3 – Training time for each classification method using the complete training dataset.

| Method | One-step Training (min) | Hierarchical classifier Training (min) | | |
|---|---|---|---|---|
| | | Sleep vs. Wake | NREM vs. REM | Total |
| Logistic Regression | 1.305017 | 0.56646388 | 0.5664639 | 1.132928 |
| Linear Discriminant Analysis | 0.325585 | 0.44011573 | 0.4401157 | 0.880231 |
| K-nearest Neighbors Classifier | 0.494233 | 0.64531681 | 0.6453168 | 1.290634 |
| Decision Tree Classifier | 3.087765 | 3.36282204 | 3.362822 | 6.725644 |
| Gaussian Naïve-Bayes | 0.169616 | 0.2236391 | 0.2236391 | 0.447278 |
| Passive Aggressive Classifier | 0.330701 | 0.25370451 | 0.2537045 | 0.507409 |
| Ridge Classifier | 0.215765 | 0.28431216 | 0.2843122 | 0.568624 |
| Logistic Regression with cross-validation | 2.72975 | 1.11243985 | 1.1124398 | 2.22488 |
| Bernoulli Naïve Bayes | 0.194801 | 0.23899159 | 0.2389916 | 0.477983 |
| Nearest Centroid | 0.138609 | 0.19050883 | 0.1905088 | 0.381018 |
| Random Forest Classifier | 2.204762 | 2.46243883 | 2.4624388 | 4.924878 |
| Ada Boost Classifier | 13.14571 | 15.7994703 | 15.79947 | 31.59894 |
| Bagging Classifier with decision tree | 19.74422 | 20.9569206 | 20.956921 | 41.91384 |
| Extra Trees Classifier | 0.650872 | 0.77603496 | 0.776035 | 1.55207 |
| Gradient Boosting Classifier | 50.63355 | 19.0567781 | 19.056778 | 38.11356 |
| Perceptron | 0.409127 | 0.22129073 | 0.2212907 | 0.442581 |
| **Bagging Classifier with Random Forest Classifier** | **14.21601** | **13.9431089** | **13.943109** | **27.88622** |
| Bagging Classifier with Extra Trees Classifier | 4.14548 | 4.19125078 | 4.1912508 | 8.382502 |

Supplementary Table 4 – Scoring time of the BCRF algorithm on subsets of the training and scoring data.

| Database | Classifier | Sleep Stages | Training Time | Scoring Time (24 hours) | Training Time with 2 hours of manual score | Scoring Time of 24 hours with 2 hours of manual score |
|---|---|---|---|---|---|---|
| **WT** | One-Step | Awake / NREM/REM | 0.8150 | 0.0086 | | |
| **DS** | One-Step | Awake / NREM/REM | 0.3515 | 0.0108 | - | - |
| **WT+DS** | One-Step | Awake / NREM/REM | 1.5338 | 0.0127 | - | - |
| **WT** | Hierarchical | Sleep/Awake | 0.9320 | 0.0107 | 0.8998 | 0.0089 |
| | | NREM/REM | 0.9651 | 0.0082 | 0.9894 | 0.0089 |
| | | Total | 1.8972 | 0.0189 | 1.8892 | 0.0178 |
| **DS** | Hierarchical | Sleep/Awake | 0.2839 | 0.0085 | 0.2905 | 0.0087 |
| | | NREM/REM | 0.2979 | 0.0074 | 0.3332 | 0.0074 |
| | | Total | 0.5818 | 0.0159 | 0.6236 | 0.0160 |
| **WT+DS** | Hierarchical | Sleep/Awake | 1.4357 | 0.0092 | 1.4296 | 0.0080 |
| | | NREM/REM | 1.6556 | 0.0083 | 1.5358 | 0.0076 |
| | | Total | 3.0914 | 0.0175 | 2.9654 | 0.0156 |

*Note.* Runtimes are listed in minutes.

Supplementary table 5 – SIESTA performance using data binned in 5-second epochs.

#### One-Step Approach

| Using complete database to score WT mice | | Using complete database to score DS mice | |
|---|---|---|---|
| **Awake** | 0.94 | **Awake** | 0.91 |
| **NREM** | 0.90 | **NREM** | 0.87 |
| **REM** | 0.75 | **REM** | 0.83 |

#### Hierarchical Approach

| Using complete database to score WT mice | | Using complete database to score DS mice | |
|---|---|---|---|
| **Awake** | 0.93 | **Awake** | 0.93 |
| **Sleep** | 0.91 | **Sleep** | 0.91 |
| **NREM** | 0.91 | **NREM** | 0.84 |
| **REM** | 0.76 | **REM** | 0.85 |

*Note.* These tests use the complete dataset (WT n=14 , DS n=6) to score one mouse of each genotype.

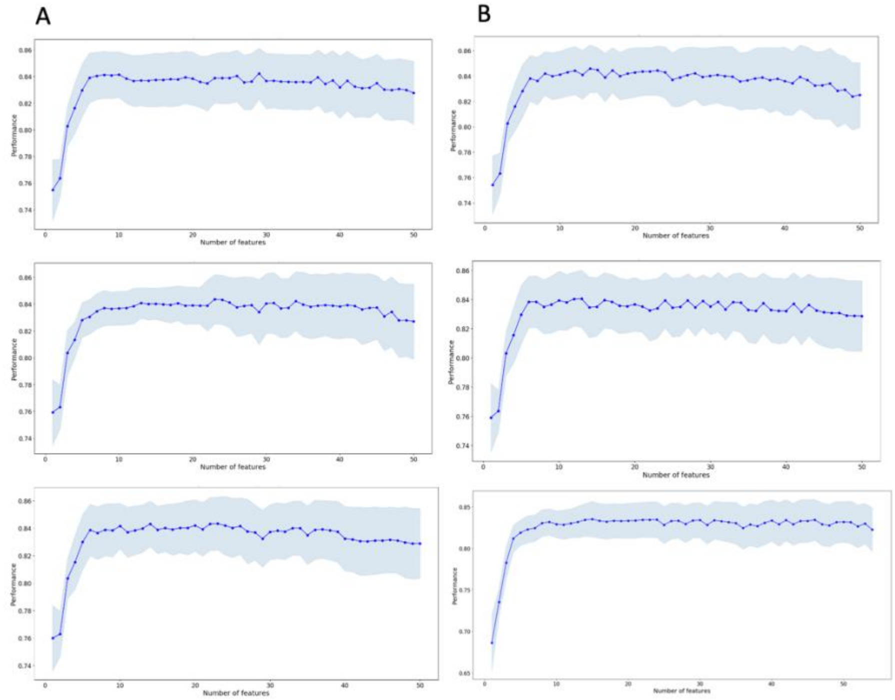


C

| Sequential Forward Selection | | |
|---|---|---|
| ECoG_zerocross | ECoG_zerocross | ECoG_zerocross |
| ECoG_spindelhan | EMG_amplitude | ECoG_spindelHan |
| EMG_amplitude | ECoG_spindelhan | ECoG_spectral_mean |
| ECoG_delta | ECoG_delta | ECoG_delta |
| EMG_spectral_entropy | EMG_spectral_entropy | EMG_sepctral_entropy |
| ECoG_thetacon | ECoG_beta | ECoG_theta3 |
| ECoG_spectral_edge | EMG_signal_var | ECoG_spectral_edge |
| ECoG_deltaenergy | ECoG_spectral_edge | ECoG_theta2 |

D

| Sequential Backward Selection. | | |
|---|---|---|
| ECoG_zerocross | ECoG_zerocross | ECoG_zerocross |
| EMG_amplitude | ECoG_spindelhan | EMG_amplitude |
| ECoG_spindelhan | ECoG_theta3amp | ECoG_spindelhan |
| ECoG_delta | EMG_spectral_entropy | ECoG_delta |
| EMG_spectral_entropy | ECoG_rms | ECoG_theta3amp |
| ECoG_theta2 | ECoG_sigma | ECoG_theta2 |
| ECoG_betaenergy | ECoG_theta2amp | ECoG_theta3 |
| ECoG_sigmaenergy | ECoG_signal_var | ECoG_sigmaamp |

**Supplementary Figure 1 –** Sequential feature selection test using the BCRF algorithm trained on the complete dataset, with a One-Step approach. The blue dots are the mean performance and the light blue shaded area is the STD in each case. **A)** Three repetitions of the Sequential Forward Selection. **B**) Three repetitions of the Sequential Backward Selection. **C)** Table with the top 8 features of each run of the Sequential Forward Selection in order of importance. **D**) Table with the top 8 features of each run of the Sequential Backward Selection in order of importance in C and D. Features highlighted in green occurred in 2 or more runs of any Sequential Selection Algorithm.

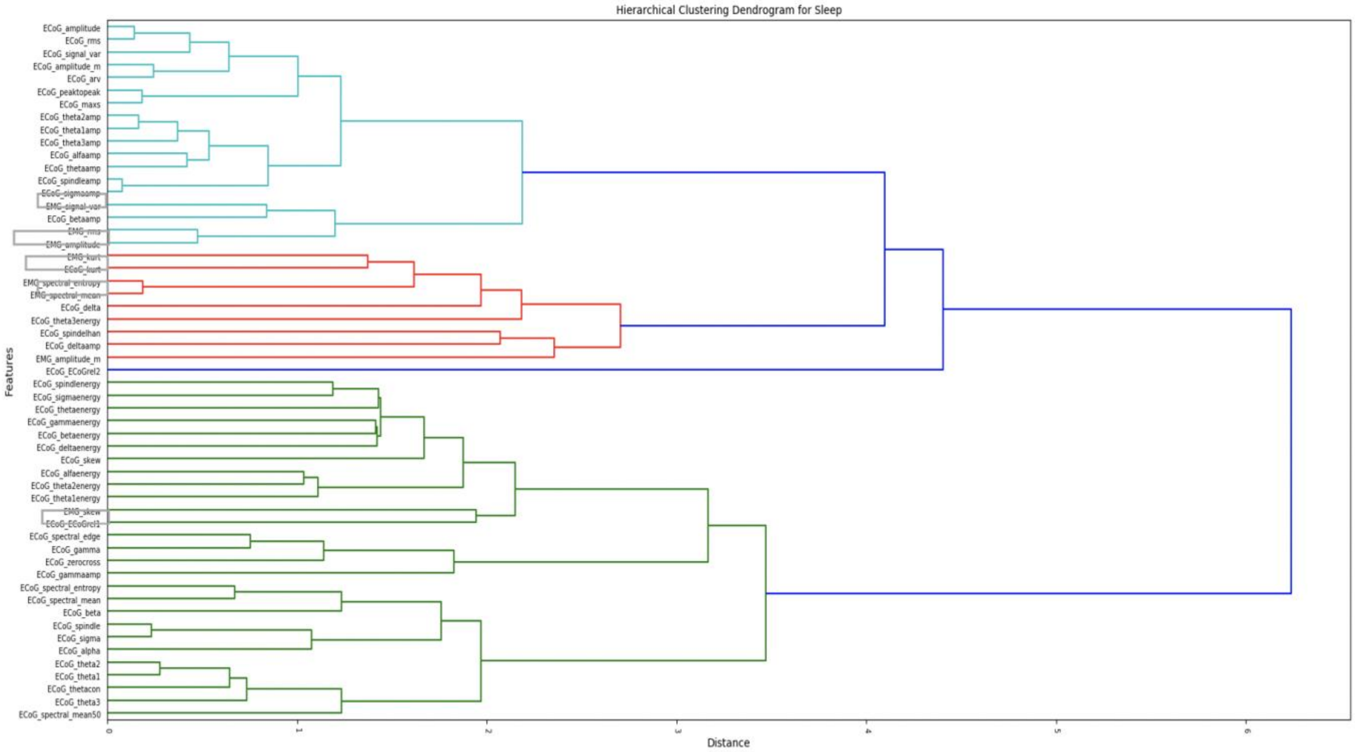


**Supplementary Figure 2 –** Cluster dendrogram of the pair-wise distance of the features from the complete dataset. In light blue, red and light green are the branches of the identified clusters. In blue the distance branches of the cluster features over the threshold. The most important features identified by SFS and SBS are boxed in grey on the y-axis.

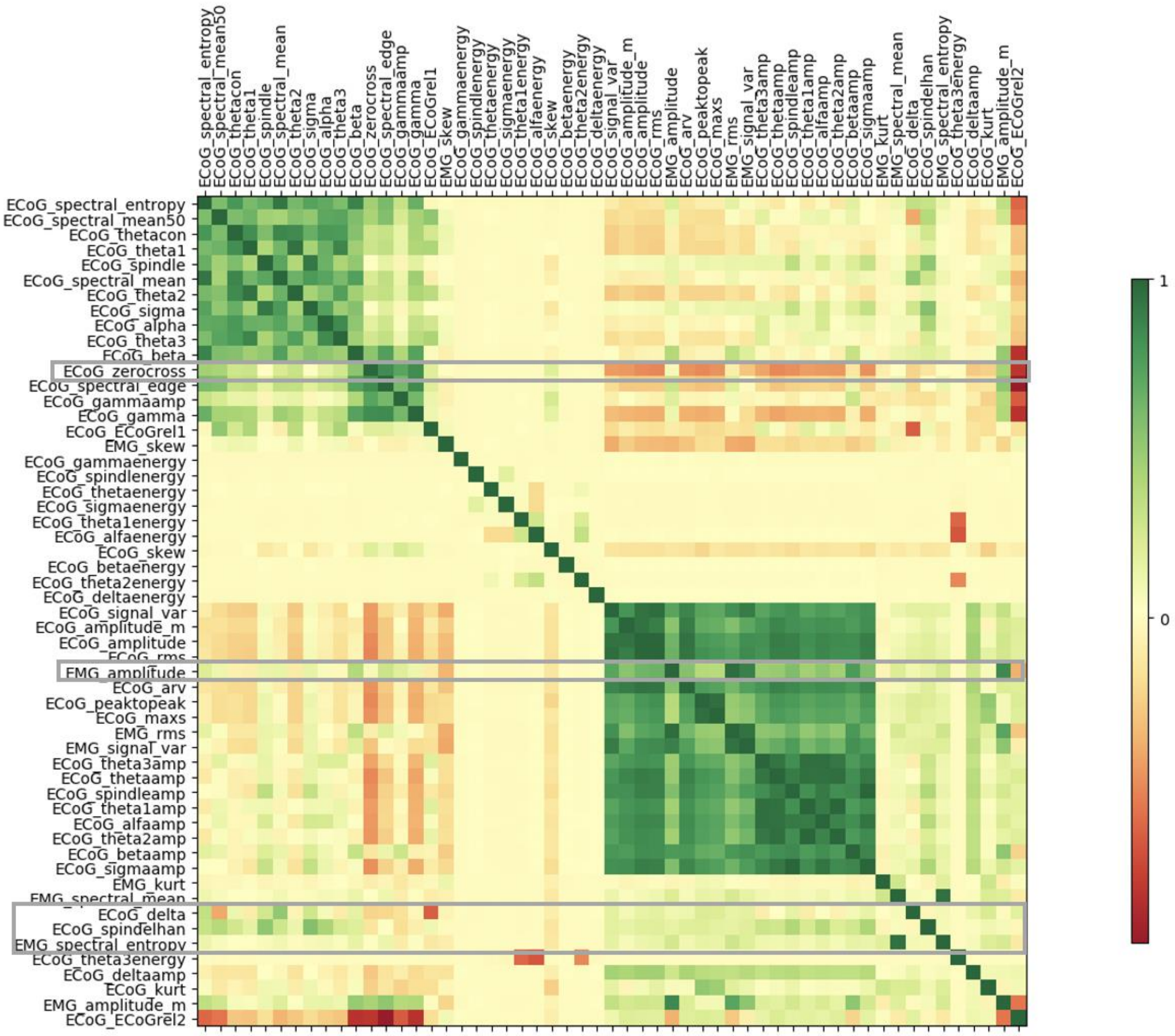


**Supplementary Figure 3 –** Same as Figure 1 in primary manuscript, with gray boxes surrounding the features consistently identified by sequential feature selection as being critical for classification performance.
